# Predicting functional consequences of recent natural selection in Britain

**DOI:** 10.1101/2023.10.16.562549

**Authors:** Lin Poyraz, Laura L. Colbran, Iain Mathieson

## Abstract

Ancient DNA can directly reveal the contribution of natural selection to human genomic variation. However, while the analysis of ancient DNA has been successful at identifying genomic signals of selection, inferring the phenotypic consequences of that selection has been more difficult. Most trait-associated variants are non-coding, so we expect that a large proportion of the phenotypic effects of selection will also act through non-coding variation. Since we cannot measure gene expression directly in ancient individuals, we used an approach (*Joint-Tissue Imputation*; JTI) developed to predict gene expression from genotype data. We tested for changes in the predicted expression of 17,384 protein coding genes over a time transect of 4500 years using 91 present-day and 616 ancient individuals from Britain. We identified 28 genes at seven genomic loci with significant (FDR < 0.05) changes in predicted expression levels in this time period. We compared the results from our transcriptome-wide scan to a genome-wide scan based on estimating per-SNP selection coefficients from time series data. At five previously identified loci, our approach allowed us to highlight small numbers of genes with evidence for significant shifts in expression from peaks that in some cases span tens of genes. At two novel loci (*SLC44A5* and *NUP85*), we identify selection on gene expression not captured by scans based on genomic signatures of selection. Finally we show how classical selection statistics (iHS and SDS) can be combined with JTI models to incorporate functional information into scans that use present-day data alone. These results demonstrate the potential of this type of information to explore both the causes and consequences of natural selection.

## Introduction

Ancient DNA (aDNA) time-series can provide direct evidence of natural selection on specific variants, avoiding confounding factors associated with inferring selection using modern data (Dehasque et al., 2020; Marciniak and Perry, 2017; Mathieson, 2020). However, in itself ancient DNA does not provide any information about the functional consequences of selection, limiting our ability to learn about phenotypes under selection and to identify effects of selection that may for example affect disease risk.

One problem is that, similar to genome-wide association studies, selection signals often span multiple genes due to linkage disequilibrium (LD), making it difficult to identify the loci targeted by selection. Indeed, long haplotypes due to selective sweeps make this problem even more challenging. One approach that has been promising in the GWAS context is to try to incorporate functional information, for example expression quantitative loci (eQTL), which have been used to link significant GWAS hits to functional consequences in transcriptome-wide association studies (TWAS) (Wainberg et al., 2019). Similarly, while the results of genome-wide association studies can be used to link signals of selection to phenotypes, without information about the intermediate functional changes, it is difficult to interpret these links.

Changes in gene expression are expected to underlie many complex traits relevant to recent human evolution (Corradin et al., 2016), particularly as many signals overlap non-coding regions of the genome. We previously used predictive models of gene expression to detect changes between different ancient subsistence groups (Colbran et al., 2021) and to infer selection based on differences between present-day populations (Colbran et al., 2023). Here, we develop this idea to test for selection directly using changes in predicted expression over time inferred from ancient DNA times series. We used an approach (*Joint Tissue Imputation* (JTI); Zhou et al., 2020) developed to predict gene expression from genotype data to predict the expression levels of ∼ 17,000 protein-coding genes in ancient (4500-1000 BP) and modern individuals from Britain. We then inferred significant shifts in expression levels in this 4500 year time-transect based on linear regression models of predicted gene expression against time.

This approach allows us to perform a gene-level test for selection on gene expression, identifying four novel signals of selection on gene expression resulting from small shifts in allele frequency that were not captured by genome-wide scans for selection. We are also able, for regions identified to have been under selection in this or other analyses, to identify which genes are likely to have changed their expression due to this selection—in several cases showing that selection at known loci (*LCT*, for example) has substantially affected the expression of several nearby genes. Our work demonstrates the utility in incorporating functional information relevant to specific hypotheses into genome-wide scans for selection.

## Results

### Imputed data recovers genome-wide selection scan results

We assembled a dataset of 91 present-day and 616 ancient (4500-1000 BP) individuals from Britain. Our approach assumes that the sample population is closed and homogeneous. We thus chose this population from Britain due to its small geographical spread, relatively continuous demographic history, and large aDNA sample size. Modern individuals were from the GBR population of the 1000 Genomes project (Auton et al., 2015). Ancient individuals had either been genotyped using the 1240k capture reagent, or shotgun sequenced and then genotyped at 1240k sites. This is the same dataset used in Mathieson and Terhorst (2022) (original sources Brace et al., 2019; Margaryan et al., 2020; Martiniano et al., 2016; Olalde et al., 2018; Patterson et al., 2022; Schiffels et al., 2016) with additional individuals from Gretzinger et al. (2022), and removing individuals with less than 0.1x coverage at 1240K sites. We calculated genotype likelihoods at 1240k sites, and then imputed diploid genotypes at 1240k sites using *beagle4* (Browning and Browning, 2007). We then lifted over 1240k sites from hg19 to hg38 and imputed at ungenotyped sites using the NHLBI TOPMed imputation server (Das et al., 2016; Fuchsberger et al., 2015; Taliun et al., 2021).

As a point of comparison to the transcriptome-wide scan, we ran the SNP-based genome-wide selection scan described in Mathieson and Terhorst (2022) on the imputed diploid 1240k dataset. Briefly, this uses the *bmws* software to estimate time varying selection coefficients based on the time series of allele frequencies, and reports P-values based on fitting a gamma distribution to the root mean squared selection coefficient, averaged in 20-SNP sliding windows. The results are largely consistent with those of Mathieson and Terhorst (2022), identifying strong evidence of selection at *LCT, DHCR7, SLC22A4, OAS1*, the HLA region and other loci (Figure 1A). Despite the larger sample size and diploid (as opposed to pseudohaploid) coverage, the new analysis does not find substantially more signals of selection and the shared signals are not more significant. Although imputation of ancient DNA generally produces accurate genotype calls (Ausmees et al., 2022; Hui et al., 2020; Sousa da Mota et al., 2023), we noticed that at strongly selected sites, imputed allele frequencies were slightly biased towards present-day allele frequencies compared to pseudohaploid allele frequencies (Supplementary Figure 1). This suggests that, while generally accurate, imputation might reduce power to detect selection because this bias has a greater effect on sites with large changes in frequency over time. In our case, this seems to offset the advantage from greater sample size. That said, the imputed data do not seem to perform worse than the pseudohaploid data, and produce well-calibrated results (Figure 1B). As imputation is necessary to perform the transcriptome-wide scan, we proceeded with the imputed dataset.

**Figure 1.**
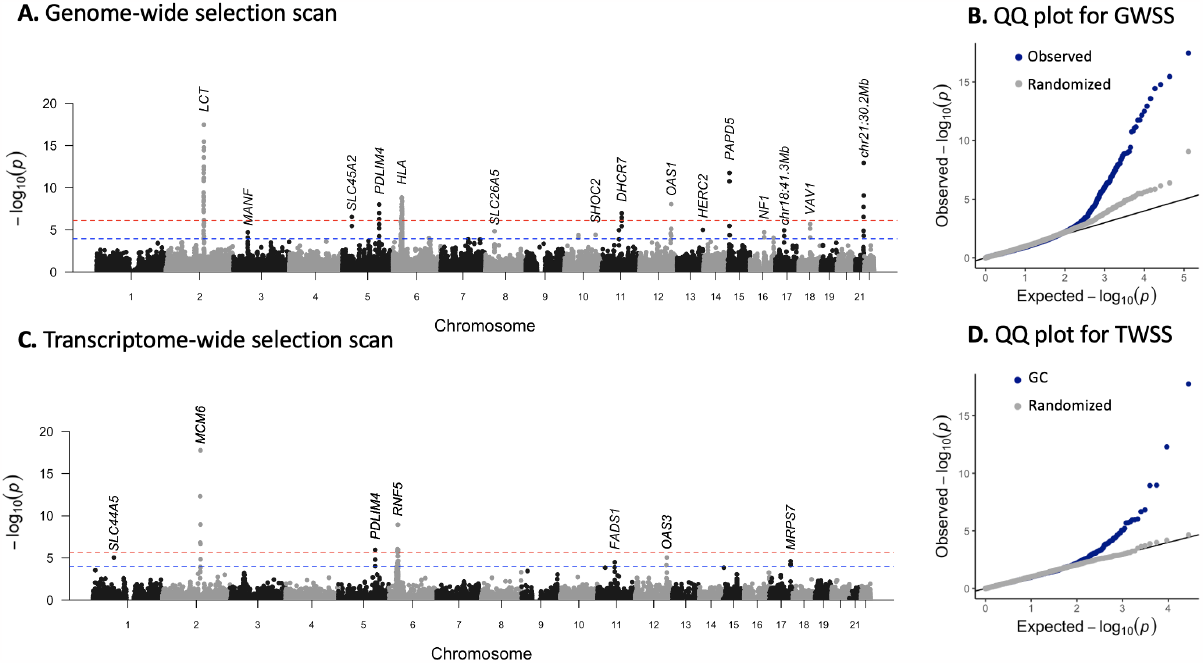
Genome-wide and transcriptome-wide scans for selection. *(A)* P-values for genome-wide selection. Each point represents a 20-SNP window. The red line indicates FDR significance (*P <* 10^*-*4^) and the blue line indicates Bonferroni significance (*P <* 10^*-*6^). FDR significant (*P <* 10^*-*4^) windows are labeled with the nearest genes or known target of selection. *(B)* QQ plot for genome-wide scan results in 20 SNP windows with points from *A* in blue, and results with dates of samples randomized in gray. *(C)* P-values for transcriptome-wide selection scan. Each point represents a gene. Red lines indicate FDR significance (*P <* 10^*-*4^) and blue lines indicate Bonferroni significance (*P <* 10^*-*6^). The most significant gene at each locus is labeled. *(D)* QQ plot for transcriptome-wide scan results with points from **C** in blue and results with dates of samples randomized in gray.

### Transcriptome-wide scan identifies significant changes in gene expression

We next carried out a transciptome-wide scan for selection. We predicted expression of 17,388 protein coding genes for each ancient and present-day individual using *JTI* models trained on the genotypes and transcriptomes of 49 tissues from the GTEx project (Zhou et al., 2020). As it is difficult to determine the most relevant tissue for each gene and models across tissues are generally correlated with each other, we used the model for the tissue with the highest training *R*^2^ for each individual gene (Colbran et al., 2023). For each gene, we fit ordinary linear regression models of expression against time to identify genes with non-neutral shifts in predicted expression levels. We did not include genetic ancestry principal components as covariates in this model, as the principal components of the ancient individuals clustered closely with the present-day individuals (Supplementary Figure 2). Instead, we applied genomic control to account for any inflation in test statistics due to genetic drift or residual population structure. After filtering for imputation quality, 28 genes at seven loci had evidence (FDR *<* 0.05) for significant shifts in predicted expression (Figure 1C, Table 1, Supplementary Figure 3).

**Table 1:** Genes with significant shifts in predicted expression (FDR *<* 0.05) characterized by the transcriptome-wide selection scan. *Tissue* indicates which tissue model was used. Note that this does not mean that the gene did not have significant shifts in other tissues, just that this tissue had the highest JTI training *R*^2^. *Beta* indicates the effect size of time on expression levels in the ordinary regression models. *GWSS Peak* indicates the significant (FDR *<* 0.05) genome-wide selection scan peak indicated in Figure 1 to which each gene corresponds.

| Chr | Gene | Tissue | P-value | Beta | GWSS Peak |
| --- | --- | --- | --- | --- | --- |
| 1 | *SLC44A5* | Skin Sun Exposed Lower leg | 8.827e-06 | -1.178e-04 | Novel |
| 2 | *TMEM163* | Kidney Cortex | 5.012e-13 | 1.400e-04 | *LCT* |
| 2 | *MAP3K19* | Testis | 1.490e-07 | 1.127e-04 | *LCT* |
| 2 | *LCT* | Cells EBV-transformed lymphocytes | 1.078e-09 | 8.239e-05 | *LCT* |
| 2 | *MCM6* | Esophagus Muscularis | 1.736e-18 | -1.795e-04 | *LCT* |
| 2 | *DARS* | Whole Blood | 2.137e-07 | -9.886e-05 | *LCT* |
| 2 | *CXCR4* | Adrenal Gland | 1.457e-05 | -4.051e-05 | *LCT* |
| 5 | *P4HA2* | Thyroid | 9.358e-05 | 1.027e-04 | *PDLIM4* |
| 5 | *PDLIM4* | Brain Cortex | 1.098e-06 | -4.210e-05 | *PDLIM4* |
| 5 | *SLC22A5* | Cells Cultured fibroblasts | 1.605e-05 | -1.283e-04 | *PDLIM4* |
| 6 | *PPP1R18* | Artery Aorta | 4.166e-05 | -2.423e-05 | HLA |
| 6 | *TUBB* | Cells EBV-transformed lymphocytes | 1.635e-06 | 4.196e-05 | HLA |
| 6 | *PSORS1C1* | Thyroid | 7.309e-05 | -7.907e-05 | HLA |
| 6 | *CDSN* | Skin Sun Exposed Lower leg | 4.200e-05 | 4.507e-05 | HLA |
| 6 | *CCHCR1* | Spleen | 9.258e-07 | 8.014e-05 | HLA |
| 6 | *APOM* | Testis | 1.935e-06 | -2.599e-05 | HLA |
| 6 | *C4A* | Brain Cerebellum | 2.017e-06 | -8.152e-05 | HLA |
| 6 | *ATF6B* | Brain Spinal cord cervical c-1 | 6.076e-06 | -5.952e-05 | HLA |
| 6 | *RNF5* | Colon Transverse | 1.145e-09 | -6.436e-05 | HLA |
| 6 | *PBX2* | Whole Blood | 1.059e-06 | -3.343e-05 | HLA |
| 6 | *HLA-DMA* | Cells Cultured fibroblasts | 8.493e-05 | 5.442e-05 | HLA |
| 6 | *HLA-DPA1* | Cells Cultured fibroblasts | 2.249e-05 | -1.380e-04 | HLA |
| 11 | *FADS1* | Brain Cerebellum | 3.310e-05 | 6.236e-05 | *FADS1* |
| 12 | *FAM109A* | Kidney Cortex | 6.886e-05 | -1.752e-05 | OAS |
| 12 | *OAS3* | Cells Cultured fibroblasts | 8.779e-06 | -1.264e-04 | OAS |
| 17 | *NUP85* | Brain Cerebellar Hemisphere | 7.025e-05 | 1.011e-04 | Novel |
| 17 | *GGA3* | Breast Mammary Tissue | 4.907e-05 | 3.052e-05 | Novel |
| 17 | *MRPS7* | Brain Cerebellar Hemisphere | 2.420e-05 | -3.548e-05 | Novel |

**Figure 2.**
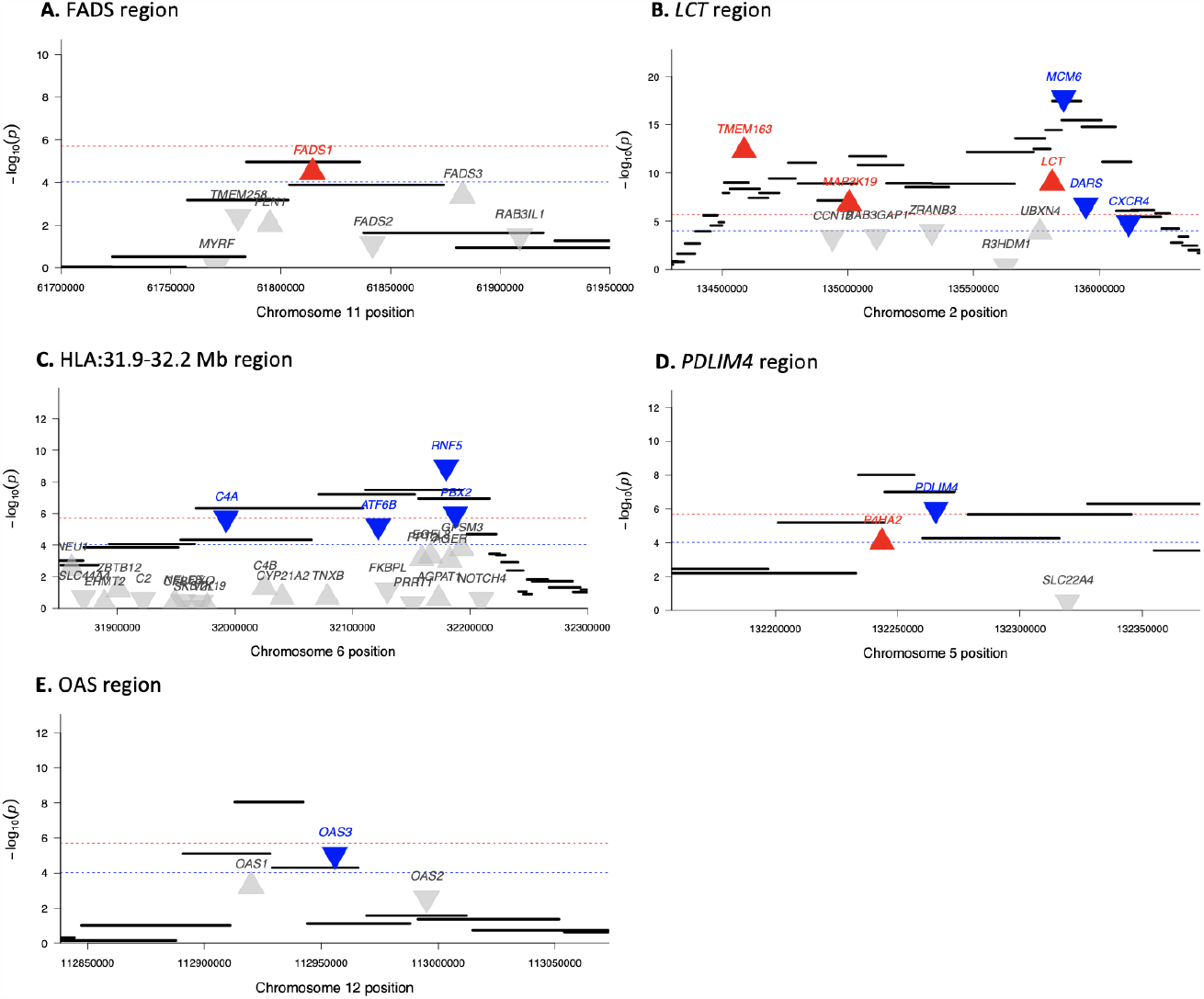
Transcriptome-wide selection scan highlights genes with directional change in expression from genomic signals of selection. Each black bar represents a 20-SNP window in the genome-wide selection scan. Each triangle indicates a gene, with up-turned and red indicating significant increased expression and down-turned and blue indicating significant decreased expression in the transcriptome-wide scan. Red lines indicate FDR significance (*P <* 10^*-*4^) and blue lines indicate Bonferroni significance (*P <* 10^*-*6^) in the transcriptome-wide scan.

**Figure 3.**
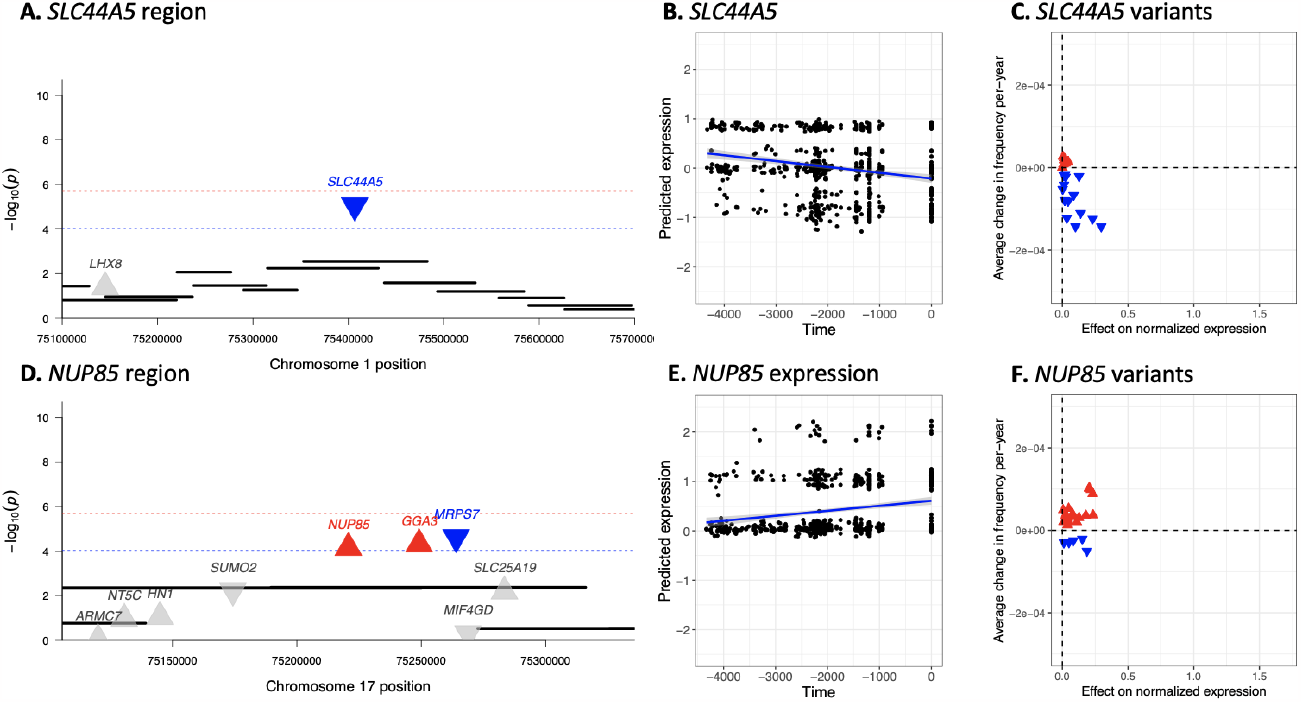
Transcriptome-wide selection scan characterizes genes with directional change in expression not captured by genome-wide selection scans. *(A,D)* Genome-wide scan for selection does not capture significant signal for selection at *SLC44A5/NUP85*. Each black bar represents a 20-SNP window in the genome-wide selection scan. Each triangle indicates a gene, with upturned and red indicating non-neutral increased expression and downturned and blue indicating non-neutral decreased expression in the transcriptome-wide scan. Red line indicates FDR significance (*P <* 10^*-*4^) and blue line indicates Bonferroni significance (*P <* 10^*-*6^) in the transcriptome-wide scan. *(B,E) SLC44A5/NUP85* expression across time. The x-axis indicates time in years before present. The y-axis indicates predicted normalized expression level. Each point represents one individual. *(C,F)* Allele frequency changes and effects of SNPs included in the prediction models for *SLC44A5/NUP85* across time. The x-axis indicates the effect of each variant on normalized expression as determined by the prediction models. The y-axis indicates average change in the frequency of each allele per year as calculated by a linear regression model of allele frequency against time. Each point represents an allele included in the prediction model for the gene. Red upturned triangles indicate alleles which have contributed to an increase in the expression level of the gene. Blue downturned triangles indicate alleles that decreased expression. Small but coordinated shifts in frequency across many alleles that were not captured by the genome-wide approach led to a decrease in the expression of *SLC44A5* and an increase in the expression of *NUP85*.

We compared the results from our transcriptome-wide scan to the SNP-based genome-wide scan. Five peaks overlap between the two scans (Figure 1). Similar to the genome-wide selection scan where peaks contain multiple SNPs (or windows of SNPs) in LD, the transcriptome-wide scan peaks span multiple genes due to both LD between eQTLs and co-regulation of nearby genes (Wainberg et al., 2019).

In some cases where the peak is shared, the transcriptome-wide scan is able to identify the genes targeted by selection as those with the largest predicted change in expression. For instance, our approach identified *FADS1* as the only gene out of the 6 in the FADS region with evidence for significant changes in expression (Figure 2B, Table 2). The genomic signal of selection in this region has been previously linked to the increased expression of *FADS1* (Ameur et al., 2012; Buckley et al., 2017; Mathieson et al., 2015), which is corroborated by our results.

**Table 2:** Genome-wide selection signal peaks and associated genes. *GWSS Start* and *GWSS End* indicate the selection peaks in the genome-wide scan for selection. Consecutive 20-SNP windows with less than 5 Mb distance in-between were merged into single signals. Three separate signals in the HLA region were merged to one signal. 0.1 Mb buffers were added to each selection signal to include all relevant genes. *TWSS significant genes* indicates the genes within the selection signal that have evidence for non-neutral regulatory shifts in the transcriptome-wide selection scan. *# All Genes* indicates the number of all protein coding genes within the selection signal, *# All Modeled Genes* indicates the number of all protein coding genes within the selection signal that were included in the TWSS after filtering for imputation quality *# Significant Genes* indicates the number of genes that achieved significance in the transcriptome-wide scan.

| Chr | GWSS Start | GWSS End | TWSS Significant Genes | # All Modeled Genes | # All Genes | # Significant Genes |
| --- | --- | --- | --- | --- | --- | --- |
| 2 | 134292811 | 136385296 | *MCM6, LCT, DARS, CXCR4, TMEM163, MAP3K19* | 11 | 12 | 6 |
| 3 | 50849763 | 51387494 |  | 1 | 1 | 0 |
| 5 | 33777346 | 34069589 |  | 2 | 4 | 0 |
| 5 | 132101060 | 132454548 | *P4HA2, PDLIM4, SLC22A5* | 5 | 5 | 3 |
| 6 | 28140757 | 33178885 | *PPP1R18, TUBB, HLA-DMA, PSORS1C1, APOM, HLA-DPA1, PBX2, RNF5, APOM, CCHCR1, CDSN, ATF6B, C4A* | 148 | 148 | 12 |
| 6 | 128796160 | 129062458 |  | 0 | 0 | 0 |
| 8 | 33218070 | 33438568 |  | 1 | 1 | 0 |
| 10 | 49643164 | 50348209 |  | 5 | 7 | 0 |
| 10 | 110778270 | 111086977 |  | 4 | 4 | 0 |
| 11 | 61684455 | 61903876 | *FADS1* | 6 | 6 | 1 |
| 11 | 71302258 | 71592390 |  | 4 | 7 | 0 |
| 12 | 110796571 | 113044151 | *OAS3, FAM109A* | 17 | 21 | 2 |
| 13 | 111558732 | 111786334 |  | 0 | 0 | 0 |
| 15 | 27951279 | 29045218 |  | 1 | 5 | 0 |
| 16 | 49972683 | 50325572 |  | 2 | 4 | 0 |
| 16 | 82983756 | 83190020 |  | 0 | 0 | 0 |
| 17 | 30984555 | 31423638 |  | 4 | 4 | 0 |
| 18 | 41362995 | 41666874 |  | 0 | 0 | 0 |
| 21 | 43286434 | 44295140 |  | 11 | 12 | 0 |

In other cases, the transcriptome-wide scan does not uniquely identify the targeted gene, but highlights a subset of genes in the region with significant changes. For example, at the *LCT* locus (Figure 2A, Table 2) the transcriptome-wide scan identified 6 out of 11 genes with significant changes in expression. As expected, we predicted a significant increase in *LCT* expression (Figure 2A), but we also predicted significant changes in five other genes. Indeed, the regulatory variants associated with adult *LCT* expression lie inside *MCM6* (Ségurel and Bon, 2017), which showed the most significant shift in expression levels in this time period.

The genome-wide scan peak at the HLA region contains 148 genes, of which we predict 12 to have significant changes in expression (Figure 2C, Table 2). The most significant signal was for *RNF5*, which is involved in the degradation of misfolded proteins and regulation of viral infection (Li et al., 2023; Zeng et al., 2021). Another signal of interest in this region is for decreased expression of *C4A*, the expression of which is associated with increased risk for schizophrenia (Yilmaz et al., 2021). These 12 genes might be priority candidates for the target of selection but it remains possible that they are all hitchhiking and the real target is a coding variant or an expression change in a gene that is not significant in our analysis.

Finally, in some cases, the gene with the most significant change in expression is probably not the main target of selection. For example, at the *PDLIM4* region (Figure 2D), we predict significant changes in expression in *PDLIM4* and *P4HA2*, but Huff et al. (2012) identified a coding variant in *SLC22A4* as the target of selection.

Similarly, although the target of selection at the OAS locus is thought to be a splice variant in *OAS1* carried by a Neanderthal introgressed haplotype, human cells with the introgressed haplotype displayed reduced *OAS3* expression and no changes in expression of *OAS1* or *OAS2* in response to viral immune triggers (Sams et al., 2016). Our transcriptome-wide scan captured this signal for reduced expression in *OAS3*, with no significant changes in predicted *OAS1* or *OAS2* expression (Figure 2E).

### Selection on gene expression not captured by SNP-based scans

Most of the genes identified by the transcriptome-wide scan fell under selection scan peaks in the genome-wide scan. However, we identified four genes at two loci with evidence for significant regulatory shifts that did not (Figure 3), Table 1). *SLC44A5* is a member of the choline transporter-like family that is highly expressed in skin, testis and esophagus. The significant predicted change in expression is due to small coordinated shifts in frequency across many alleles (Figure 3). *SLC44A5* is one of relatively few genes with a population-biased eQTL (GTEx Consortium, 2020). Specifically, rs4606268 has a much larger effect on *SLC44A5* expression in European ancestry individuals compared to those of African ancestry, consistent with rapid evolution of the regulation of this gene in European populations. *SLC44A5* is also more highly expressed in lymphoblastoid cell lines (LCLs) of European ancestry, compared to African ancestry. However British (GBR) LCLs show the lowest expression (Supplementary Figure 4; Lappalainen et al., 2013), consistent with our finding of recent selection for lower expression in Britain. Positive selection at *SLC44A5* has previously been reported in East Asian populations (Yasumizu et al., 2020) and JTI predictions suggest low expression levels (Supplementary Figure 4). Since *SLC44A5* is highly expressed in skin, and skin pigmentation experienced strong selection in both Britain and East Asia, we hypothesize that selection on *SLC44A5* expression may also be related to skin pigmentation. Choline is closely related to folate, which is broken down by UV radiation and thought to drive selection for darker skin pigmentation in high-UV regions (Jablonski and Chaplin, 2010). In mice, choline partially counteracts the effects of low folate in development (Craciunescu et al., 2010), so one possibility is that selection on *SLC44A5* acts to counteract the increased rate of folate degradation due to light skin pigmentation.

**Figure 4.**
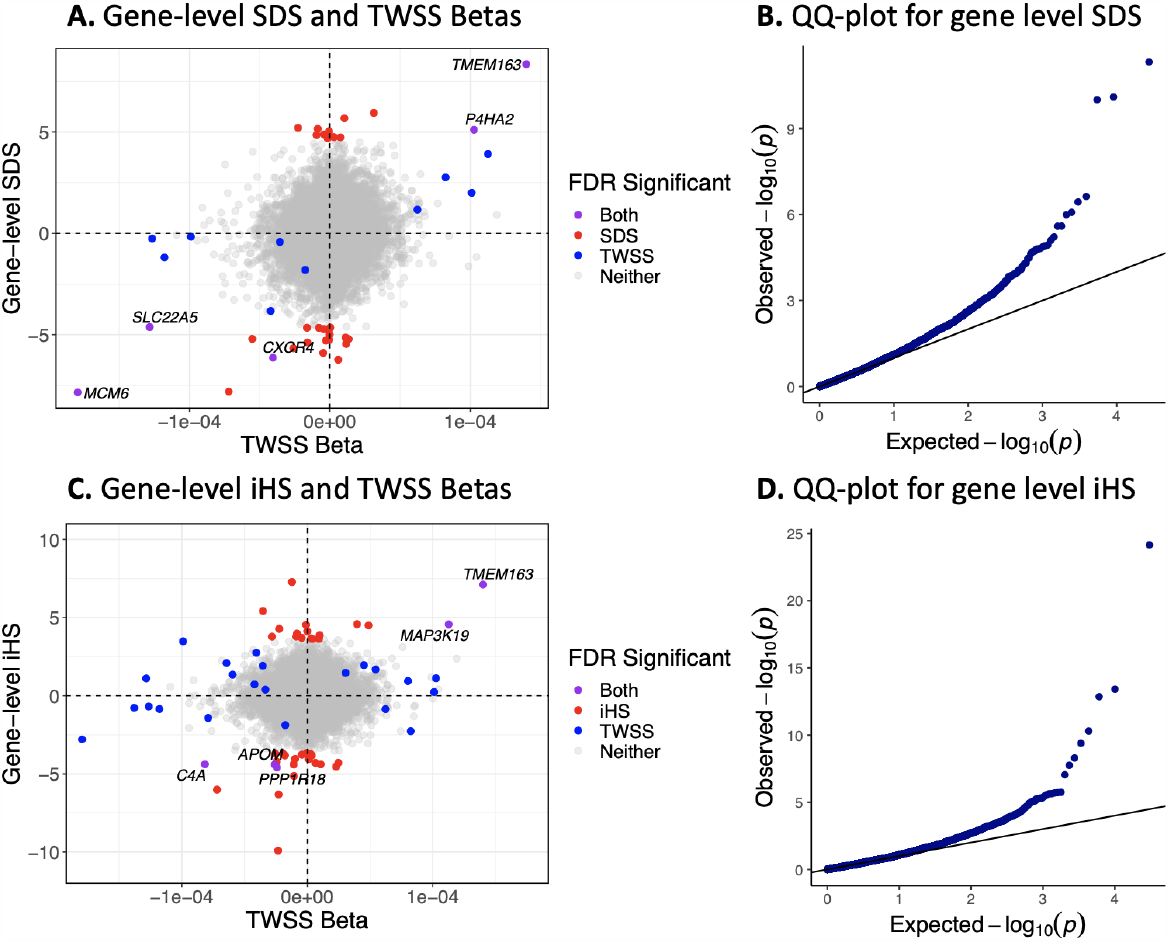
iHS and SDS statistics combined with functional information reveal the gene-level consequences of selection. *(A,C)* Gene-level SDS/iHS and average change in predicted gene expression per year. The x-axis indicates average change in predicted gene expression per year as measured by the beta of time in the transcriptome-wide selection scan and the y-axis indicates gene-level SDS/iHS values. Positive gene-level scores indicate an increase in gene expression resulting from selection, whereas negative scores indicate a decrease in expression. Each point represents a gene, with blue points indicating genes that reached FDR significance in the transcriptome-wide scan, red points indicating those that reached significance in the gene-level SDS/iHS analysis scan, gray points indicating genes that do not reach FDR significance in either scan, and purple indicating genes that achieve significance in both. There is greater concordance between the SDS analysis and the transcriptome-wide scan compared to the iHS scan, as expected by the similar time-scales of the SDS and the transcriptome wide scan analyses. *(B,D)* QQ plot for gene-level SDS/iHS with genomic control imposed.

The second novel locus includes *NUP85, GGA3*, and *MRPS7*. It is likely that one of these genes is the target of selection, as all protein-coding genes within 100kb of this signal were modeled. Of these three, *NUP85* is upregulated in GBR LCLs compared to the other European 1kG populations, as would be expected under our inferred scenario of selection for increased expression on this gene in Britain (Supplementary Figure 4). *NUP85* encodes a part of the nucleoporin complex, which controls transport between the cytoplasm and the nucleus (Ling et al., 2022). It is involved in the recruitment and migration of immune cells through chemokine signaling (Toda et al., 2009), as well as the control of viral replication (Brass et al., 2008; Ling et al., 2022), suggesting pathogen-induced selective pressures.

### Classical selection statics can also be combined with functional information

Although ancient DNA provides direct evidence of selection, its usefulness is limited by sample size, data quality and limited geographic and temporal availability. We therefore also explored a complementary approach of combining the JTI models with classical selection statistics based on present-day populations. To do this, we generated gene-level selection statistics from SNP-level selection statistics based on the integrated haplotype score (iHS) and the singleton density score (SDS) (Field et al., 2016; Voight et al., 2006). The test statistic is a standardized weighted sum of the per-SNP selection statistics included in the predictive model for each gene weighted by effect size on normalized gene expression. As both SDS and iHS are already normalized, this weighted sum also has a standard normal distribution. We therefore calculated P-values based on Z scores, and applied genomic control to account for residual correlation between JTI model SNPs. SDS and iHS were not available for each SNP included in our prediction models, so we removed genes for which less than half of the SNPs had scores available.

Based on SDS scores, 34 genes had significant evidence for selection (FDR *<* 0.05). Five of these (*MCM6, TMEM163, P4HA2, CXCR4*, and *SLC22A5*) were significant in both the gene-level SDS and the transcriptome-wide selection scan. 13 genes that were significant in the transcriptome-wide scan were filtered out of the gene-level SDS analysis due to missing SDS scores. Based on iHS scores, 48 genes had FDR *<* 0.05, of which 5 were also significant in the transcriptome-wide scan (*TMEM163, MAP3K19, C4A, APOM*, and *PPP1R18*). 1 gene that was significant in the transcriptome-wide scan was filtered out of the gene-level iHS analysis due to missing iHS scores.

Given that the SDS scan detects selection in the last ∼ 2000 years, we expected it to be more correlated with the transcriptome-wide scan compared to iHS which captures the last ∼ 10000 years. In terms of shared significiant signals, 5/34 is not significantly different than 5/48. However, we note that all 15 genes that achieved significance in the transcriptome-wide scan and had gene-level SDS available had the same predicted direction of change in gene expression in the SDS analysis. On the other hand, the predicted direction of changes in gene expression in the iHS analysis were uncorrelated with those predicted by the transcriptome-wide scan (*ρ* = 0.0438).

These results show that functional information can be used to enhance selection scans based on standard selection statistics, as well as tests specifically designed to use the eQTL data (Colbran et al., 2023). However, as with SNP-based scans, the information obtained from these statistics is complementary to information obtained from ancient DNA time series. While ancient DNA allows direct observation of selection and precise estimates of timing, present-day samples can be much larger and therefore more powerful, though potentially more sensitive to artefacts and model misspecification.

## Discussion

In this study, we used JTI models to detect selection on gene expression over the last 4500 years in Britain. We identified 28 genes (FDR *<* 0.05) with evidence for selection, of which 24 were identified by a SNP-based genome-wide selection scan on the same data. Importantly, the transcriptome-wide scan identified significant shifts in predicted expression of genes that were not captured by scans based on SNP-based genomic signatures of selection that do not incorporate functional information. Though we focused here on the application to ancient DNA data, we also demonstrated how eQTL data can be incorporated into selection scans based on present-day data, with complementary results.

The results of the transcriptome-wide scan can be interpreted in multiple ways. First, where significant genes overlap with peaks from SNP-based genome-wide scans, the transcriptome-wide scan can be used to prioritize genes that are targets of selection. This is analogous to the way in which eQTL colocalization is helpful but not a complete solution to identifying causal genes at genome-wide association peaks. The most significant gene may not be the most important, or may not have a JTI model, or the target of selection may be a coding variant (e.g. at *SLC45A2*, where we find no significant genes in the transcriptome-wide scan). Nonetheless, we find several examples where the most significant gene in the transcriptome-wide scan is the targeted gene at a genome-wide scan peak. Second, the transcriptome-wide scan can identify genes (for example *SLC44A5*) that are not identified in the genome-wide scan because the selection is relatively polygenic. Finally, the transcriptome-wide scan can identify the effects of selection on genes that are not themselves the target of selection. For example, selection on the expression of *LCT* affects the expression of several other genes which may themselves have functional consequences. More generally, although the interpretation of selection scans tends to focus on a single causal gene at a locus, the transcriptome-wide scan makes it clear that the linked and co-regulated genes can be important and the phenotypic changes that selection acts on reflect the composite effect of many genes. For example, we predict that selection on lactase persistence significantly changed the expression of at least five other genes and the fitness consequences of the selected allele would depend on the aggregate effects of these changes.

Our approach still has several technical limitations. First, it is tissue-agnostic. Because expression is typically highly correlated across tissues, we focused on the tissue with the highest *R*^2^ in our scan. However, the tissue with the highest expression is not necessarily the one that is the target of selection. While appropriate for a general scan, more tissue-specific predictions can be used to test specific hypotheses (such as melanocytes in the case of selection on skin pigmentation (Colbran et al., 2021)) Second, even for most genes, the JTI models explain only a relatively small proportion of the variance in expression and include only *cis*-regulatory variants. Many genes cannot be modeled at all, and eQTLs themselves may be context-specific in which case we could miss signals of selection entirely.

Overall, this study demonstrates the potential of incorporating functional predictive models in the analysis of ancient DNA to explore the phenotypic drivers and consequences of selection. Our transcriptome-wide scan for selection provides a broad overview of the regulatory shifts associated with recent human evolution in Britain, and shows how the TWAS workflow can be used to better understand the molecular basis and consequences of selection.

## Methods

### Data collection and imputation

We identified ancient individuals from Britain with genome-wide ancient DNA data, dated to within the past 4500 years (Brace et al., 2019; Gretzinger et al., 2022; Margaryan et al., 2020; Martiniano et al., 2016; Olalde et al., 2018; Patterson et al., 2022; Schiffels et al., 2016). Most of these data had been generated using the 1240k capture reagent but some had been shotgun sequenced. We calculated genotype likelihoods at 1240k sites using a binomial model for read counts with a 1% error rate and a 5% deamination rate. We then imputed diploid genotypes at 1240k sites using *beagle4* (Browning and Browning, 2007) with the 1000 Genomes reference panel (Auton et al., 2015). We then lifted over the 1240k sites from hg19 to hg38, and imputed ungenotyped sites using the NIH TOPMed server (Das et al., 2016; Fuchsberger et al., 2015; Taliun et al., 2021). Finally, we merged these genotypes with present-day individuals data from the GBR population of the 1000 Genomes Project Phase 3 NYGC resequenced data using *bcftools* (Byrska-Bishop et al., 2022; Danecek et al., 2021). We removed genetic ancestry PCA outliers and individuals with less than 0.1x coverage at 1240K sites, retaining a total of 616 ancient and 91 present-day individuals.

### Genome-wide selection scan

We ran a genome-wide scan for selection based on selection coefficient estimation from time series aDNA data as described in Mathieson and Terhorst (2022). We used the imputed data at 1240k sites, lifted over to hg38. We started with 1,150,639 autosomal SNPs and filtered out all SNPs with MAF *<* 0.1, greater than 90% missingness and those with MAF = 0 in the ancient data leaving 409,232 SNPs. We inferred selection coefficients at each generation using a smoothing parameter, λ = 10^4.5^ and effective population size *N*_*e*_ = 10^4^. We calculated root mean squared selection coefficients for 20-SNP sliding windows sliding in 10-SNP increments. We fit a gamma distribution to the window selection coefficients and computed P-values for each window.

### Models for predicted gene expression

In order to construct predictive models for gene expression, we used published JTI gene expression models, which leverage shared regulation across tissues (Zhou et al., 2020). These models were trained on common variants (MAF *>* 0.05) for 49 tissues in version 8 of the Genotype Tissue Expression project (GTEx) (GTEx Consortium, 2020). For each gene, we utilized the tissue with the highest training *R*^2^ as described by Colbran et al. (2023). The median number of SNPs in each model was 12. The median *R*^2^ for these models was 0.1938.

### Transcriptome-wide selection scan

We constructed ordinary linear regression models of predicted expression against time for 17,833 protein-coding genes:

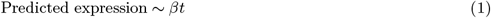

where t indicates years before present and *β* indicates average change in predicted expression per year. We did not include genetic ancestry principal components as covariates in this model, as the principal components of the ancient individuals clustered closely with the modern individuals. We calculated imputation quality scores for each gene 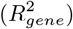 by taking a weighted average of the quality scores of each SNP included in the prediction model for each gene 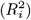 with weights |*β*_*i*_| equal to the absolute JTI effect size of the SNP on normalized gene expression:

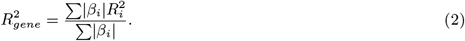

We filtered out the 20% genes with the lowest imputation quality, retaining 13,892 genes. We applied genomic control to the resulting p-values to account for genetic drift. We calculated an inflation factor,, *λ* = 1.791, and divided all test statistics by, *λ* to ensure that the median p-value was equal to the median p-value in the null *χ*^2^ distribution (Devlin and Roeder, 1999).

We also randomized the dates of the samples and re-ran the linear regression analysis to generate randomized p-values. We categorized genes that achieved FDR significance (P *<*0.0001) as those with evidence for significant changes in predicted expression.

### Gene-level iHS and SDS analysis

We generated gene-level selection statistics from SNP-level classical selection statistics, the integrated haplotype score (iHS) and the singleton density score (SDS) (Field et al., 2016; Voight et al., 2006). We retrieved SDS scores calculated using data from 3195 individuals from Britain in the UK10K dataset from Field et al. (2016). For the iHS analysis, we used data from 91 individuals from the GBR population in the 1000 Genomes Project (Auton et al., 2015). We polarized the ancestral/derived alleles of the 1000G individuals with respect to the chimpanzee reference genome (GenBank accession: GCA 002880755.3). We then used selscan with the -norm flag to calculate normalized iHS scores (Szpiech and Hernandez, 2014).

We calculated gene-level selection statistics by taking the sum of the SDS/iHS values of each SNP included in the predictive model for each gene multiplied by its effect size on normalized gene expression (*β*_*i*_). SDS and iHS were not available for every SNP included in our prediction models, so we removed genes for which less than half of the SNPs had scores available. We retained a total of 13,588 genes for the SDS analysis, and 15,123 genes for the iHs analysis. As the SNP-level SDS and iHS scores were normalized, we re-normalized the gene-level selection scores by dividing by square root of the sum of the squared effect sizes:

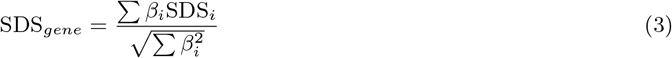

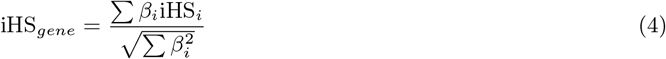

where the sums are over all SNPs *i* in the model for each gene. We treated the normalized gene-level statistics as z-scores to generate p-values, to which we applied genomic control (Devlin and Roeder, 1999).

## Data and Code Availability

All data used are publicly available from original sources. Summary statistics for the transcriptome-wide scan are included as supplementary table 1. Code for generating figures and running analyses is available at: https://github.com/linpoyraz/predicting_functional_britain.

