## Supplementary Figures 1-3 for "Predicting functional consequences of recent natural selection in Britain"

### Supplementary Materials

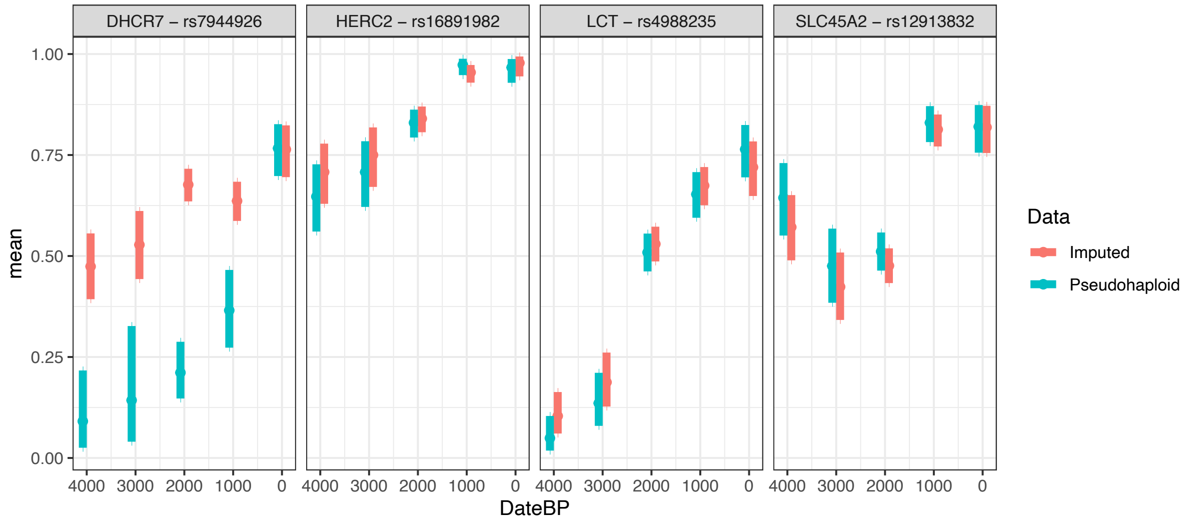


Supplementary Figure 1: Estimated derived allele frequencies and 95% confidence intervals for samples in 1000-year bins for four different strongly selected variants. Pseudohaploid data in blue and imputed dipliod data in red. In the first three cases, the imputed allele frequencies are biased towards the present-day frequencies, suggesting that selection signals at these loci would be attenuated.

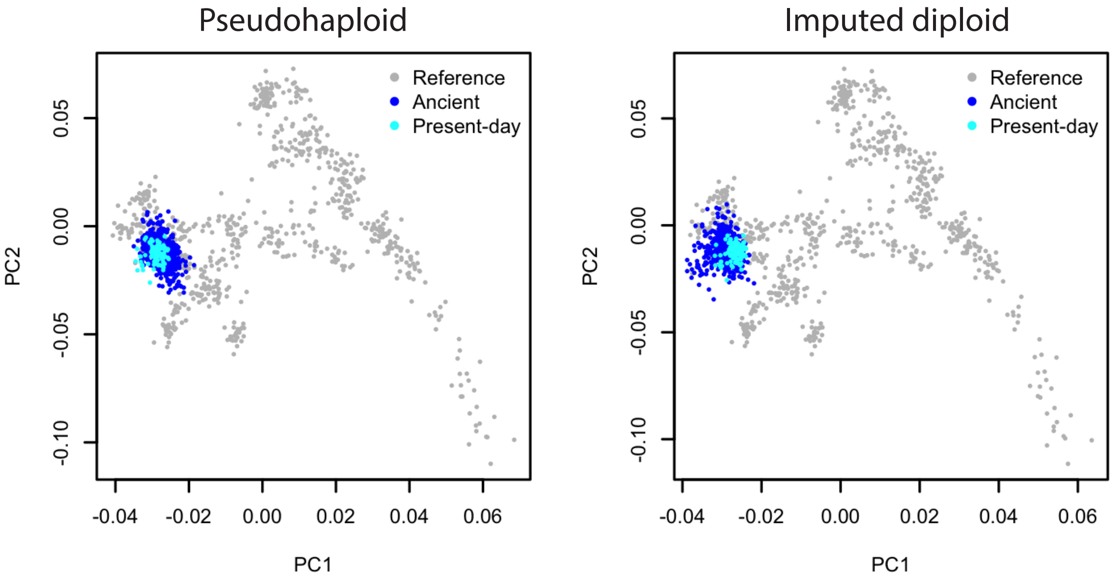


Supplementary Figure 2: PCA plots of ancient (dark blue) and present-day (light blue) samples used in the analysis projected onto principal components defined by 777 present-day West Eurasian samples genotyped on the Human Origins array (see Lazaridis et al. (2014) for details and population labels). In the left panel the ancient samples are pseudohaploid, while in the right panel they are imputed to diploid coverage. Only sites included on the Human Origins array are included.

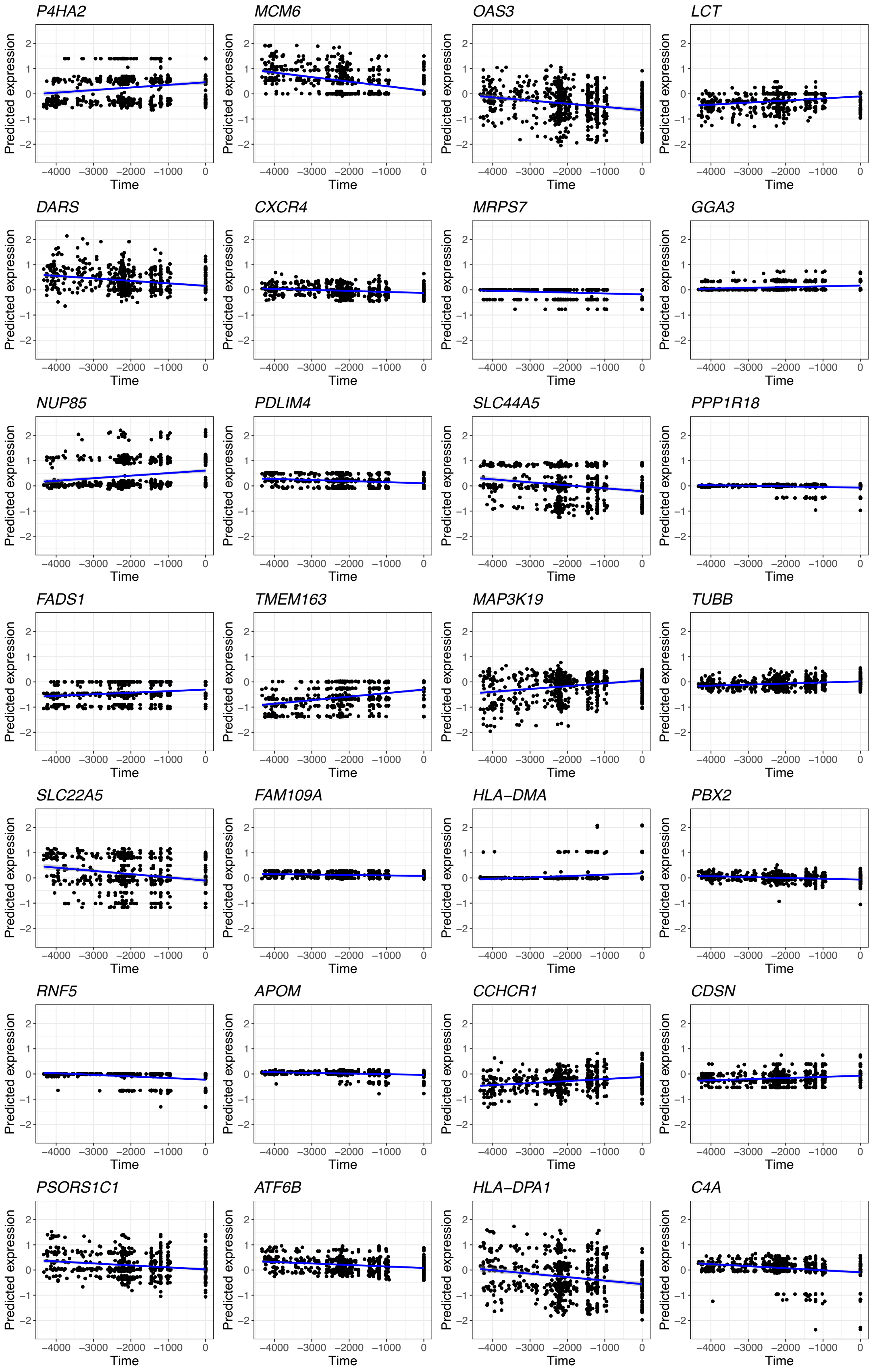


Supplementary Figure 3: Changes in gene expression across time for significant genes. The x-axis indicates time, where 0 represents modern samples. The y-axis indicates predicted normalized expression level. Each point represents a sample.

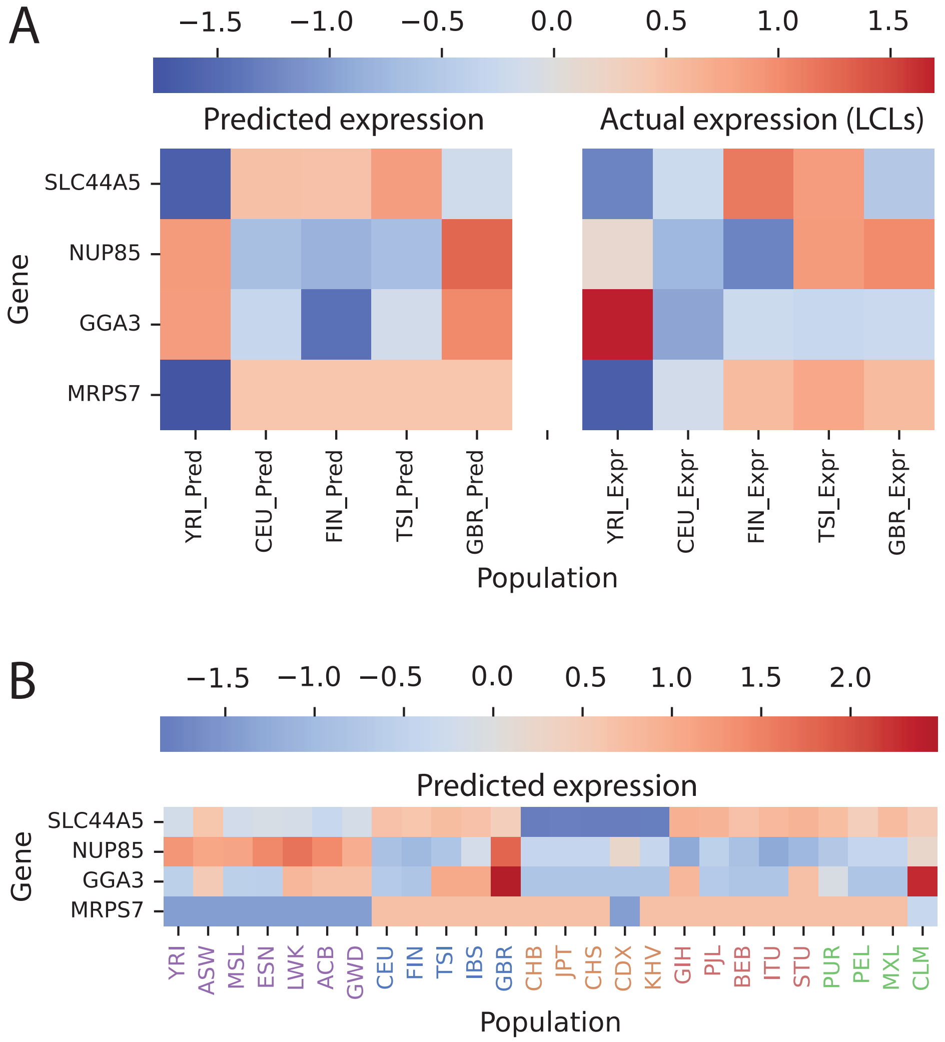


Supplementary Figure 4: Predicted and actual expression of genes at novel signals. Heatmaps show values normalised so that the standard deviation over individuals equals 1. **A**: Median predicted expression based on JTI models (Zhou et al., 2020) and median actual expression in LCLs (Lappalainen et al., 2013) for one African (YRI) and four European (CEU, TSI, FIN, GBR) populations. **B**: Median predicted expression based on JTI models for all 1000 Genomes Populations (Auton et al., 2015).
